# Transcription elongation factor ELOF1 is required for efficient somatic hypermutation and class switch recombination

**DOI:** 10.1101/2024.09.24.614732

**Authors:** Lizhen Wu, Anurupa Devi Yadavalli, Gabriel Matos-Rodrigues, Dijin Xu, Andreas P. Pintado-Urbanc, Matthew D. Simon, Wei Wu, André Nussenzweig, David G. Schatz

## Abstract

Somatic hypermutation (SHM) and class switch recombination (CSR) diversify immunoglobulin (Ig) genes and are initiated by the activation induced deaminase (AID), a single-stranded DNA cytidine deaminase that is thought to engage its substrate in the context of RNA polymerase II (RNAPII) transcription. Through a loss of function genetic screen, we identified numerous potential factors involved in SHM including ELOF1, a component of the RNAPII elongation complex that has been shown to function in DNA repair and transcription elongation. Loss of ELOF1 strongly compromises SHM, CSR, and AID targeting and alters RNAPII transcription by reducing RNAPII pausing downstream of transcription start sites and levels of serine 5 but not serine 2 phosphorylated RNAPII throughout transcribed genes. ELOF1 must bind to RNAPII to be a proximity partner for AID and to function in SHM and CSR. We propose that ELOF1 helps create the appropriate stalled RNAPII substrate on which AID acts.

**Highlights:**

- A CRISPR knockout screen has identified numerous potential SHM factors.
- SHM, CSR, and AID targeting are strongly compromised in the absence of ELOF1.
- ELOF1 must interact with RNAPII to be an AID proximity partner and support AID targeting.
- ELOF1 supports RNAPII pausing and generation of the substrate for AID action.

## Introduction

Ig genes are diversified in antigen-activated B cells by SHM and CSR to facilitate effective humoral immune responses. SHM introduces point mutations into Ig heavy and light chain variable regions to enable the generation of high affinity antibodies while CSR involves DNA double strand break intermediates in Ig heavy chain switch regions and replaces one heavy chain constant region with another to alter antibody effector function. Both reactions are initiated by the activation induced deaminase (AID), a deoxycytidine deaminase that acts specifically on single-stranded (ss) DNA substrates to create deoxyuridine, processing of which yields SHM and CSR products.^1–3^ SHM and CSR require RNAPII transcription, which is thought to be the source of the ssDNA on which AID acts.^4^ Components of the RNAPII elongation complex have been implicated in SHM and/or CSR, some as AID interaction partners,^5–11^ and several lines of evidence suggest a link between transcription pausing/stalling and AID targeting.^3, 4, 6, 12–14^. In particular, recent evidence suggests that the SHM targeting function of Ig enhancers is mediated by their ability to increase RNAPII stalling in SHM target regions.^15^ However, it remains unclear in what context or how AID gains access to ssDNA since the transcription bubble is buried within paused and elongating RNAPII complexes.^16–18^ CSR can be induced to occur at high efficiency in as little as two days in immortalized cell line and *ex vivo* primary B cell culture systems, enabling successful genetic screens for CSR factors.^6, 19^ The ability to perform genetic screens for SHM factors has been hampered by the lack of an equivalent rapid and sensitive assay system. To overcome this, we recently developed RASH (<u>R</u>apid <u>A</u>ssay for <u>S</u>omatic <u>H</u>ypermutation), a system based on Ramos human Burkitt Lymphoma B cell lines engineered to contain a sensitized GFP-based SHM reporter integrated into the *Ig heavy chain* (*IGH*) locus and inducible expression of AID7.3.,^20^ a catalytically hyperactive form of AID.^21^ In the RASH-1 cell line, SHM can be assessed after two to four days of AID induction by loss of either GFP fluorescence or surface IgM expression using flow cytometry, or by DNA sequencing. We have now performed a CRISPR-Cas9 loss of function screen in RASH-1, resulting in the identification of a number of novel potential SHM factors, including ELOF1 (elongation factor 1 homologue).

ELOF1 is a small (83 aa), evolutionarily conserved, constitutive component of elongating RNAPII.^17, 22, 23^ Structural studies demonstrate that human ELOF1 and its yeast orthologue Elf1 bind directly to RNAPII to form part of the DNA entry channel.^17, 24^ ELOF1/Elf1 help orchestrate transcription-coupled nucleotide excision repair (TC-NER)^22–25^ and have been implicated in facilitating transcription through nucleosomes and protecting against transcription-induced DNA replication stress in cells with high levels of transcription blocking DNA lesions.^22, 23, 26, 27^ Disruption of *ELOF1* in human adherent cell lines leads to a high sensitivity to UV irradiation due to defective TC-NER but few reported defects in the absence of DNA damage other than a reduced RNAPII elongation rate.^22, 23^ Here, we demonstrate that ELOF1 participates in three-way complexes with AID and RNAPII and contributes significantly to both SHM and CSR in a manner that requires its interaction with RNAPII. Our results identify a new function for ELOF1, link ELOF1 to RNAPII pausing and AID targeting, and implicate the balance between RNAPII pausing and elongation as a critical control point for AID function.

## Results

### Loss of function SHM screen identifies numerous factors related to RNAPII transcription

A genome-wide CRISPR-Cas9 loss of function screen in RASH-1 cells (Figure 1A) identified numerous potential factors involved in SHM, with *AICDA* (the gene encoding AID) as the second ranked positive hit (sgRNAs enriched in GFP positive cells) (Table S1). Gene ontology analysis of the top 200 positive hits showed strong enrichment for genes associated with RNAPII activity (Figure S1A). To validate the screen, we expressed individual sgRNAs targeting 33 of the top 200 hits in bulk cultures of RASH-1C (an approach hereafter referred to as “bulk KO”). We found that about half of the hits could be validated, with higher ranked hits being validated more efficiently than lower ranked hits (Figure 1B and S1B, S1C). Validated hits included zinc finger proteins such as ZNF541, ZNF829, ZNF45, ZNF468, ZNF578, ZNF575, and transcription factors ZEB1 and CTBP1, which have been reported to form a repressive complex at a distal promoter element of *BCL6,*^28^ one of the most mutated non-Ig genes in human and mouse germinal center B cells.^29, 30^ The strongest SHM phenotype observed in this analysis was for *ELOF1* (rank 113), for which GFP loss and IgM loss decreased ∼3-fold in bulk KO experiments (Figure 1C). These findings suggest that ELOF1 participates in SHM.

**Figure 1.**
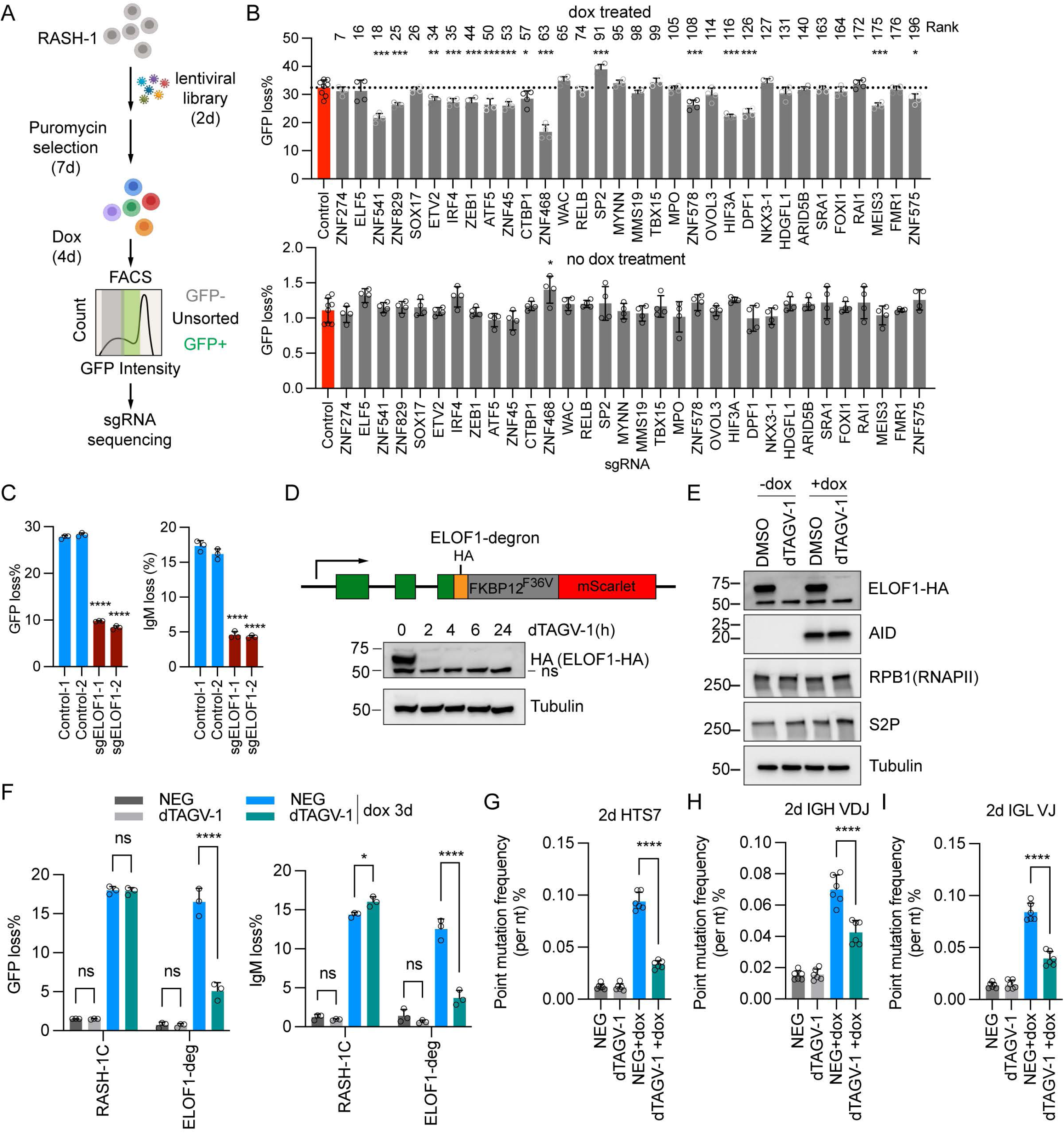
Genome-wide CRISPR screen in RASH-1 cells reveals RNAPII machinery involved in SHM. (A) Schema showing the genome-wide CRISPR screening workflow. (B) RASH-1C cells treated with empty vector control (red bars) or expressing sgRNA against CRISPR screen hits from gene ontology categories marked with red dots in Figure S1A with (top) or without (bottom) 4-day dox treatment were assayed for GFP loss. MAGeCK screen rank is indicated above the bar. (C) GFP loss and IgM loss in RASH-1C cells treated with control or two different *ELOF1* sgRNAs with 4-day dox treatment. ELOF1 ranked 113 in the screen. (D) Schematic of the edited endogenous ELOF1 gene fused with degron tag (top). Western blot in ELOF1-degron RASH-1C cells with indicated time course of dTAGV-1 treatment (bottom). (E) Western blot of ELOF1-degron RASH-1C cells with dTAGV-1 treatment for 4 hours followed by 1-day dox treatment. RPB1 is the large subunit of RNAPII. S2P, S2P-RNAPII. (F) GFP loss and IgM loss in RASH-1C or ELOF1-degron cells treated with NEG or dTAGV-1 for 4 hours followed by 3-day dox treatment. NEG, negative control for dTAGV-1. (G-I) Point mutation frequencies in *HTS7* (G), *IGH VDJ* (H) and *IGL VJ* (I) region from ELOF1-degron RASH-1C cells treated with dTAGV-1 for 4 hours followed by 2-day dox or no dox treatment. Throughout the figure, data are presented with bar representing mean and error bar as ± SD. Statistical significance was calculated using one way ANOVA with Dunnett’s post-test for B, C, G-I and two-way ANOVA with Tukey’s multiple comparisons test for F. ****p value<0.0001, ***p value<0.001; **p value<0.01; *p value<0.05.

### ELOF1 is required for efficient SHM and CSR

Despite several attempts, we were unable to obtain homozygous *ELOF1* knockout (KO) RASH-1 clones (although heterozygous KOs were readily obtained), suggesting that ELOF1 is required for long-term cell viability and/or proliferation in these cells. We therefore used the degradation tag (dTAG) system^31^ to generate RASH-1 cells in which ELOF1 could be rapidly degraded, integrating the degron cassette so that it would be expressed fused to the C-terminus of endogenous *ELOF1* (Figure 1D). dTAGV-1 treatment led to substantial degradation of ELOF1 in 2 hours and to undetectable levels of ELOF1 in 4 hours (Figures 1D and S1D). Importantly, degradation of ELOF1 for 24 hours had no detectable effect on levels of expression of RPB1 (the large subunit of RNAPII), RNAPII with its C-terminal domain (CTD) Serine 2 phosphorylated (S2P), or doxycycline-(dox)-inducible AID, and 2-day dTAGV-1 treatment had no detectable effect on cell viability or cell proliferation (Figures 1E and S1E). Consistent with the results obtained with *ELOF1* sgRNAs, we observed a ∼3-fold decrease of both GFP loss and IgM loss upon dTAGV-1 treatment as compared to cells treated with the dTAGV-1 negative control compound (NEG) (Figure 1F). In contrast, dTAGV-1 treatment did not decrease GFP or IgM loss in WT RASH-1 cells, arguing against non-specific effects of dTAGV-1 (Figure 1F). High-throughput sequencing revealed a significant dTAGV-1-dependent decrease in point mutation frequencies (Figures 1G-1I) and deletion and insertion frequencies (Figures S1F-S1K) at *HTS7*, *IGH-VDJ* and *IGL-VJ* without alterations to the mutation spectrum (Figures S1L-S1N). To exclude the possibility that the SHM defect caused by degradation of ELOF1 was related to dox-inducible hyperactive AID7.3, we generated *ELOF1*-degron WT Ramos cells. Degradation of ELOF1 in WT Ramos for 5 days did not change endogenous AID expression and had no significant effect on cell proliferation or cell viability (Figures S2A and S2B) but led to 2.5-fold decrease in IgM loss and significant decreases in point mutation frequencies in AID hotspots in both *IGH-VDJ* and *IGL-VJ* (Figures S2C-S2E).

The observation that loss of ELOF1 reduced point mutation, deletion, and insertion frequencies while leaving the mutation spectrum unaltered, prompted us to hypothesize that ELOF1 functions to facilitate the action of AID rather than in subsequent error-prone repair of AID-generated uracils. If this is the case, then degradation of ELOF1 should lead to reduced levels of uracil in SHM target regions. To test this prediction, we measured uracil abundance throughout the genome with USER-S1-END-seq,^32^ a method that employs sequential enzymatic steps (Figure 2A) to convert uracils into double-stranded DNA breaks that are then detected by END-seq, a genome-wide high-throughput sequencing method.^32, 33^ USER-S1-END-seq signal was readily detected in ELOF1-degron RASH-1 cells on both strands in *IGH-VDJ* and *IGL-VJ*, was strongly decreased in dTAGV-1 treated cells, and was completely dependent on AID induction with dox (Figures 2B and S2F). In contrast, the USER-S1-end-seq signal was not decreased by dTAGV-1 treatment in WT RASH-1C cells (Figure S2G). These results demonstrate that ELOF1 degradation reduces AID-dependent uracil levels in Ig V regions, consistent with a role for ELOF1 in facilitating AID-mediated deamination to initiate SHM.

**Figure 2.**
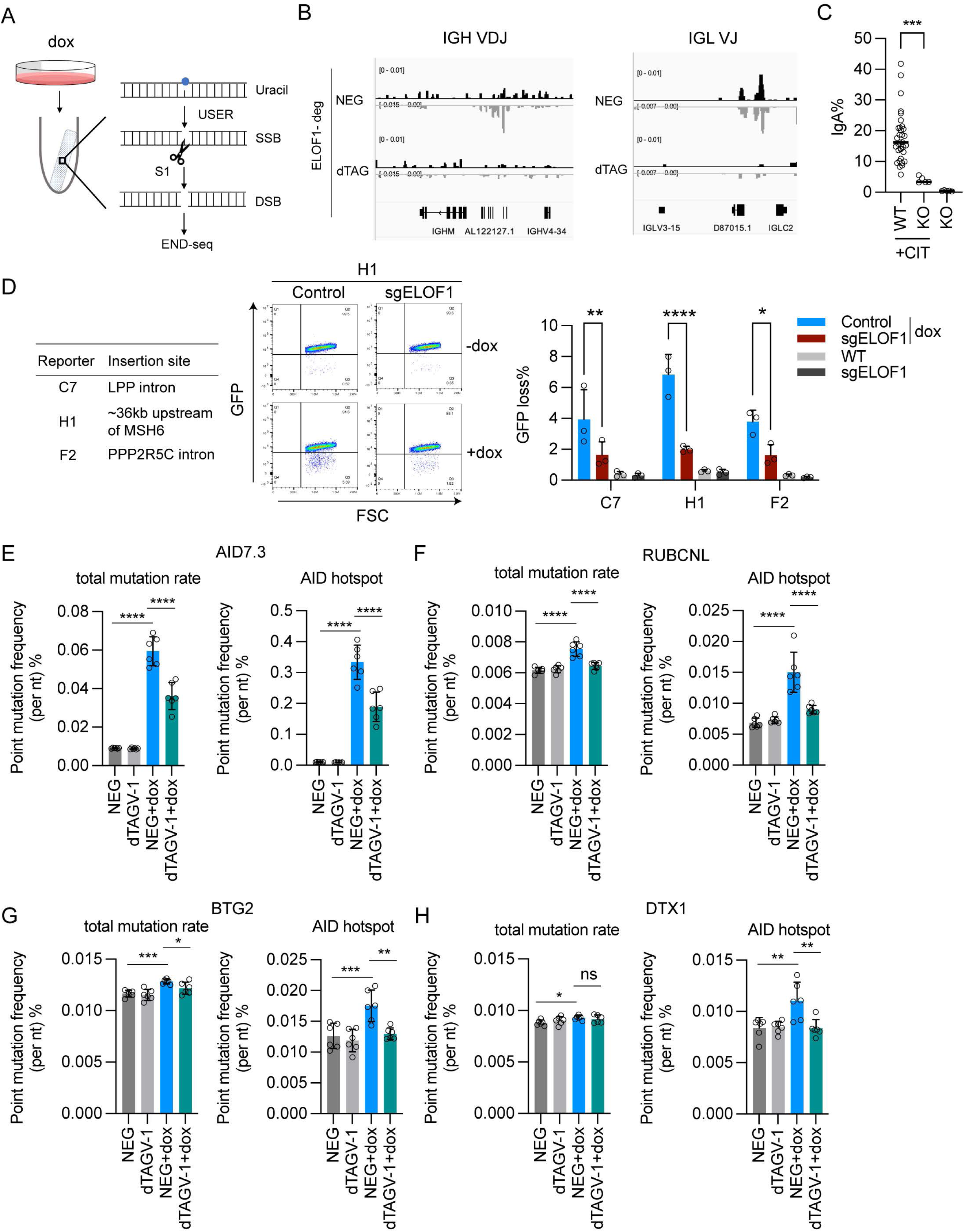
ELOF1 is required for efficient SHM and CSR and AID activity at SHM off-target loci. (A) USER-S1-END-seq experimental strategy. USER: a combination of an uracil DNA-glycosylase and endonuclease VIII. (B) USER-S1-END-seq genome browser track showing peaks in *IGL VDJ* and *IGL VJ* regions from ELOF1-degron RASH-1C cells treated with NEG or dTAGV-1 for 24 hours. (C) *ELOF1* KO CH12F3 cells were induced with CIT for 3 days and then assayed for CSR from IgM to IgA. CIT: anti-CD40, IL-4 and TGF-β1. (D) Table showing location of insertion of SHM reporter in AID off-target reporter cell lines (left). Representative flow cytometry plots of and bar graphs quantifying GFP loss from AID off-target reporter cells treated with control or *ELOF1* sgRNA induced with dox for 5 days. (E-H) Total point mutation frequency and AID hotspot point mutation frequency in *AID7.3* (E), *RUBCNL* (F), *BTG2* (G) and *DTX1* (H) regions from ELOF1-degron RASH-1C cells treated with dTAGV-1 for 4 hours followed by 5-day dox or no dox treatment. Throughout the figure, data are presented with bar representing mean and error bar as ± SD. Statistical significance was calculated using one way ANOVA with Dunnett’s post-test for C, E-H and Two-way ANOVA with Tukey’s multiple comparisons test for D. ****p value<0.0001; ***p value<0.001; **p value<0.01; ns, not significant.

We tested whether ELOF1 plays a role in CSR by knocking out ELOF1 in CH12F3 cells, which undergo cytokine-dependent inducible switching to IgA.^34^ While 17.8% of WT CH12-F3 could be induced to switch to IgA, this value was reduced 4.5-fold, to 4%, in five independent ELOF1 KO cell lines (Figure 2C). Loss of ELOF1 did not affect Sμ or Sα germline transcript levels in CIT stimulated cells (Figure S2H) or expression of AID (see below). Taken together, our results indicate that ELOF1 is required for efficient SHM and CSR.

### ELOF1 facilitates AID activity at SHM off-target loci

AID targets numerous non-Ig genes at low levels, with significant consequences for B cell oncogenesis.^29, 30, 35^ We previously demonstrated that *GFP7-E*, the lentiviral SHM reporter vector used in the creation of RASH-1, can be used to identify regions of the genome susceptible or resistant to SHM.^36^ To investigate whether ELOF1 contributes to AID off-target activity, we integrated *GFP7-E* into Ramos cells with dox-inducible AID7.3, identified single-cell clones with substantial AID-dependent GFP loss, and selected three clones for further analysis. The three clones (C7, H1, and F2) had *GFP7-E* integrated into different non-Ig regions of the genome (Figure 2D) that we had previously found to be highly susceptible to SHM.^36^ Bulk KO of *ELOF1* in these three cell lines significantly reduced GFP loss compared to cells with intact ELOF1 (Figure 2D). Hence, SHM of the reporter at several non-Ig sites in the genome is dependent on ELOF1.

We extended the analysis using high-throughput sequencing to assess mutation frequencies in ELOF1-degron RASH-1 cells at eight endogenous non-Ig loci previously identified as SHM targets in Ramos.^37^ Four of the eight loci (*AID7.3*, *RUBCNL*, *BTG2*, and *DTX1*) exhibited an increase in mutations upon AID induction, and in all four cases, degradation of ELOF1 led to a significant reduction in the mutation frequency, most clearly observed for mutations in AID hotspots (Figures 2E-2H and S2I-S2L). Taken together, our results argue that ELOF1 enables the action of AID at Ig V regions, Ig S regions, and non-Ig off-target loci.

### ELOF1 must interact with RNAPII to function in SHM and CSR

ELOF1 is composed a three domains, a positively charged N-terminus, a zinc-finger, and a C-terminal acidic tail that mediates interaction with RNAPII (Figure 3A). Double mutation of S72 and D73 to K (SDK mutant) has been shown to disrupt the ELOF1-RNAPII interaction.^23^ SDK mutant ELOF1 expressed well in RASH-1 cells while deletion of the first 15 aa or double mutation of C26 and C29 zinc coordinating residues resulted in much lower protein levels compared to wild type ELOF1 (Figure 3A). To investigate the mechanism by which ELOF1 functions in SHM, we expressed WT or SDK mutant ELOF1 in ELOF1-degron RASH-1C cells treated with dTAGV-1 or DMSO vehicle (Figures 3B-3E and S3A-S3F). Upon ELOF1 degradation, GFP loss, IgM loss, and frequencies of point mutations, deletions, and insertions could be rescued by reconstitution with WT ELOF1 but not by SDK mutant ELOF1, which yielded results that were not significantly different from empty vector in most cases (Figures 3B-3E and S3A-S3F). We performed similar reconstitution experiments in *ELOF1* KO CH12F3 cells and observed strong reconstitution of CSR with WT but not SDK mutant ELOF1 (Figure 3F). Western blotting demonstrated that SDK mutant ELOF1 was expressed at least as well as WT ELOF1 and that AID expression was similar between WT and ELOF1 KO CH12F3 cells and was not affected by ELOF1 reconstitution (Figure S3G). These results indicate that ELOF1 exerts its function in SHM and CSR through its binding to RNAPII.

**Figure 3.**
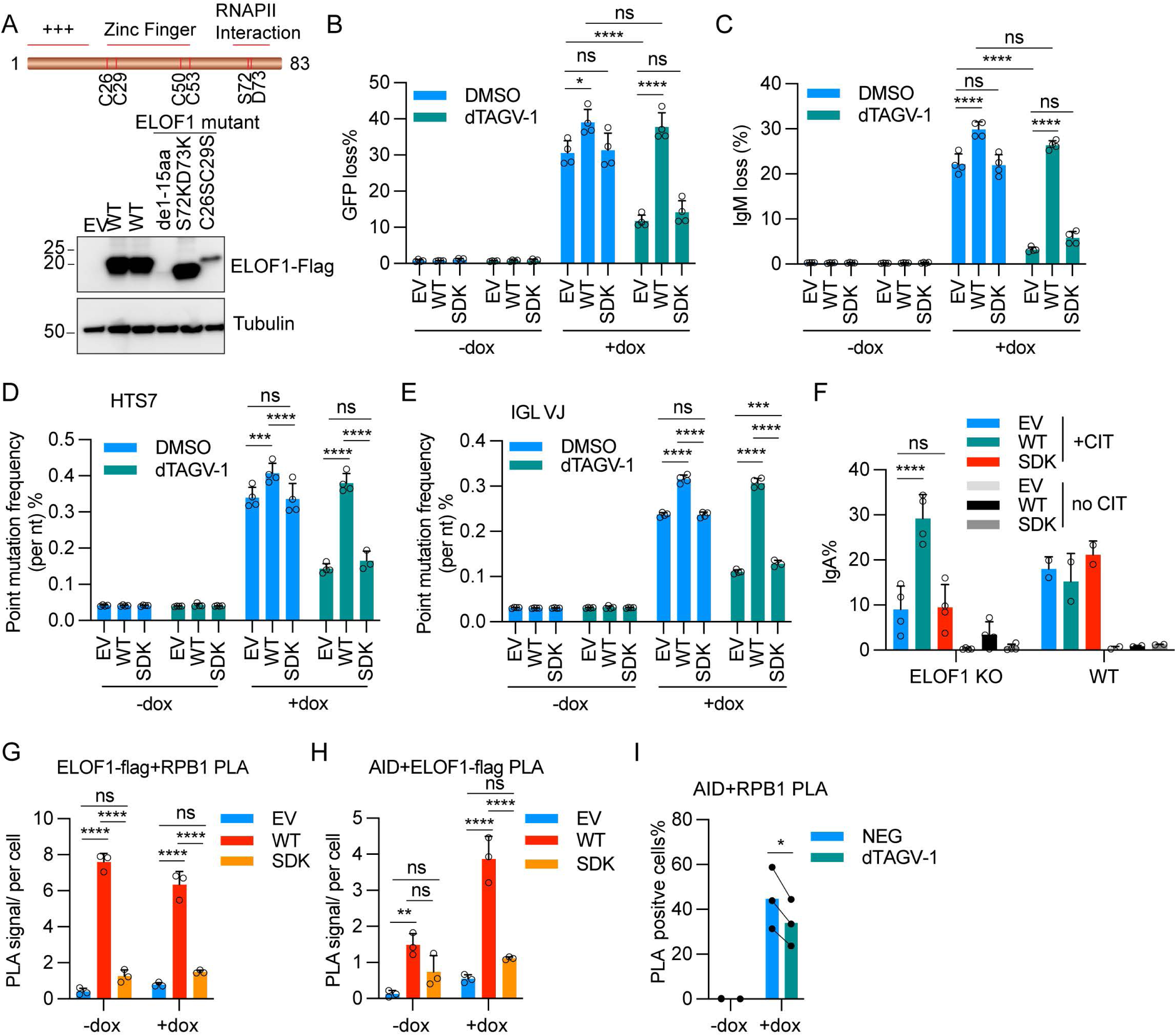
ELOF1 must interact with RNAPII to function in SHM and CSR. (A) Schematic of human ELOF1 protein (top). Western blot showing ELOF1 protein levels in RASH-1C cells overexpressing WT ELOF1 or its mutants. (B, C) WT ELOF1 or SDK mutant was expressed in ELOF1-degron RASH-1C cells. Cells treated with dTAGV-1 for 4 hours followed by 4-day dox treatment were assayed for GFP loss (B) and IgM loss (C). (D, E) Point mutation frequencies in *HTS7* (D), and *IGL VJ* (E) regions from ELOF1-degron RASH-1C cells reconstituted with WT ELOF1 or SDK mutant treated with dTAGV-1 for 4 hours followed by 4-day dox or no dox treatment. (F) CSR from IgM to IgA in *ELOF1* KO CH12F3 cells reconstituted with WT ELOF1 or SDK mutant treated with CIT for 3 days. (G, H) ELOF1-degron RASH-1C cells reconstituted with WT ELOF1 or SDK mutant treated with dTAGV-1 for 4 hours followed by 1-day dox or no dox treatment. PLA was done using flag antibody (ELOF1-flag) and RPB1 antibody in (G) or flag antibody (ELOF1-flag) and AID antibody in (H). PLA signals quantified by imaging. (I) ELOF1-degron RASH-1C cells reconstituted with WT ELOF1 or SDK mutant treated with NEG or dTAGV-1 for 4 hours followed by 1-day dox or no dox treatment. PLA was done using AID antibody and RPB1 antibody. PLA signals quantified by FACS. Throughout the figure, data are presented with bar representing mean and error bar as ± SD. Statistical significance was calculated using Two-way ANOVA with Tukey’s multiple comparisons test for B-I. ****p value<0.0001; ***p value<0.001; **p value<0.01; ns, not significant.

### Evidence for an AID-ELOF1-RNAPII complex

We tested for the existence of a three-way AID-ELOF1-RNAPII complex using the proximity ligation assay (PLA) in cells expressing WT or SDK mutant ELOF1. Consistently, we observed PLA signal between ELOF1 and RPB1 by imaging (Figure 3G) and between ELOF1 and RPB1, S2P-RNAPII, or S5P-RNAPII by flow cytometry (Figures S3H-S3J). ELOF1-RNAPII PLA signals were independent of AID expression and abrogated with SDK mutant ELOF1, confirming that the SDK mutation disrupts ELOF1-RNAPII association (Figures 3G and S3H-S3J). In addition, AID and ELOF1 proximity was readily observed in cells expressing AID and the PLA signal was lost with SDK mutant ELOF1 (Figures 3H and S3K). Hence, AID and ELOF1 are proximity partners specifically in the context RNAPII.

We then asked whether loss of ELOF1 affects the association between AID and RNAPII. Using PLA and flow cytometry, we observed an AID-RNAPII PLA signal specifically in cells expressing AID and a small (∼20%), statistically significant decrease in the signal in the absence of ELOF1 (Figure 3I). Co-immunoprecipitation (co-IP) with anti-AID antibody yielded pull down of RPB1, ELOF1, S5P-RNAPII, and SPT5 (a multifunctional RNAPII pause/elongation factor^38, 39^ that interacts with AID^6^) in a manner dependent on expression of AID (Figure S3L). The co-IP signal for RPB1, S5P-RNAPII, and SPT5 was reduced slightly or was not detectably altered upon ELOF1 degradation (Figure S3L). We conclude that ELOF1, RNAPII, and AID can exist together in a complex, that AID can associate with RNAPII in the absence of ELOF1, and that ELOF1 makes at most a small contribution to the AID-RNAPII association in the context of cells performing SHM.

### ELOF1 modulates RNAPII occupancy and pausing in AID target genes

To better understand the function of ELOF1 in SHM, we used Cleavage Under Targets & Release Using Nuclease sequencing (CUT&RUN-seq)^40^ to map ELOF1 chromatin occupancy genome-wide, which to our knowledge has not been described. Experiments were performed in ELOF1 degron RASH-1C cells treated with either NEG or dTAGV-1 and with dox for two days, taking advantage of the HA tag present on the ELOF1 degron protein. Parallel CUT&RUN-seq experiments mapped RPB1 (total RNAPII), S5P-RNAPII, S2P-RNAPII, and H3K4me3, which marks active TSS regions.^41^ The ELOF1 metagene profile showing peaks at transcription start sites (TSS) and transcription end sites (TES) and occupancy throughout the transcription unit, with the signal largely abrogated in dTAGV-1 treated cells (Figure 4A). The distribution of ELOF1 in the genome correlates particularly tightly with that of S2P-RNAPII (Figures 4B; correlation coefficient of 0.94), consistent with previous studies that found ELOF1 associated with S2P-RNAPII.^22, 23^ Notably, the H3K4me3 and S2P-RNAPII metagene profiles are unaltered by degradation of ELOF1, while total RNAPII redistributes from the TSS to the gene body and S5P-RNAPII is reduced at the TSS in the absence of ELOF1 (Figures 4C-4F). UV irradiation also results in a redistribution of total RNAPII into the gene body, an effect strongly exacerbated if ELOF1 is absent, likely due to accumulation of RNAPII at DNA lesions in gene bodies.^22, 23^ Our data indicate that loss of ELOF1 leads to a decrease in total and S5P-RNAPII at TSSs and gene body accumulation of a form of RNAPII that is not well recognized by the S5P- and S2P-RNAPII antibodies used. This pattern was maintained for genes of different length (0-50 kb, 50-100 kb, >100 kb) (Figures S4A-S4C).

**Figure 4.**
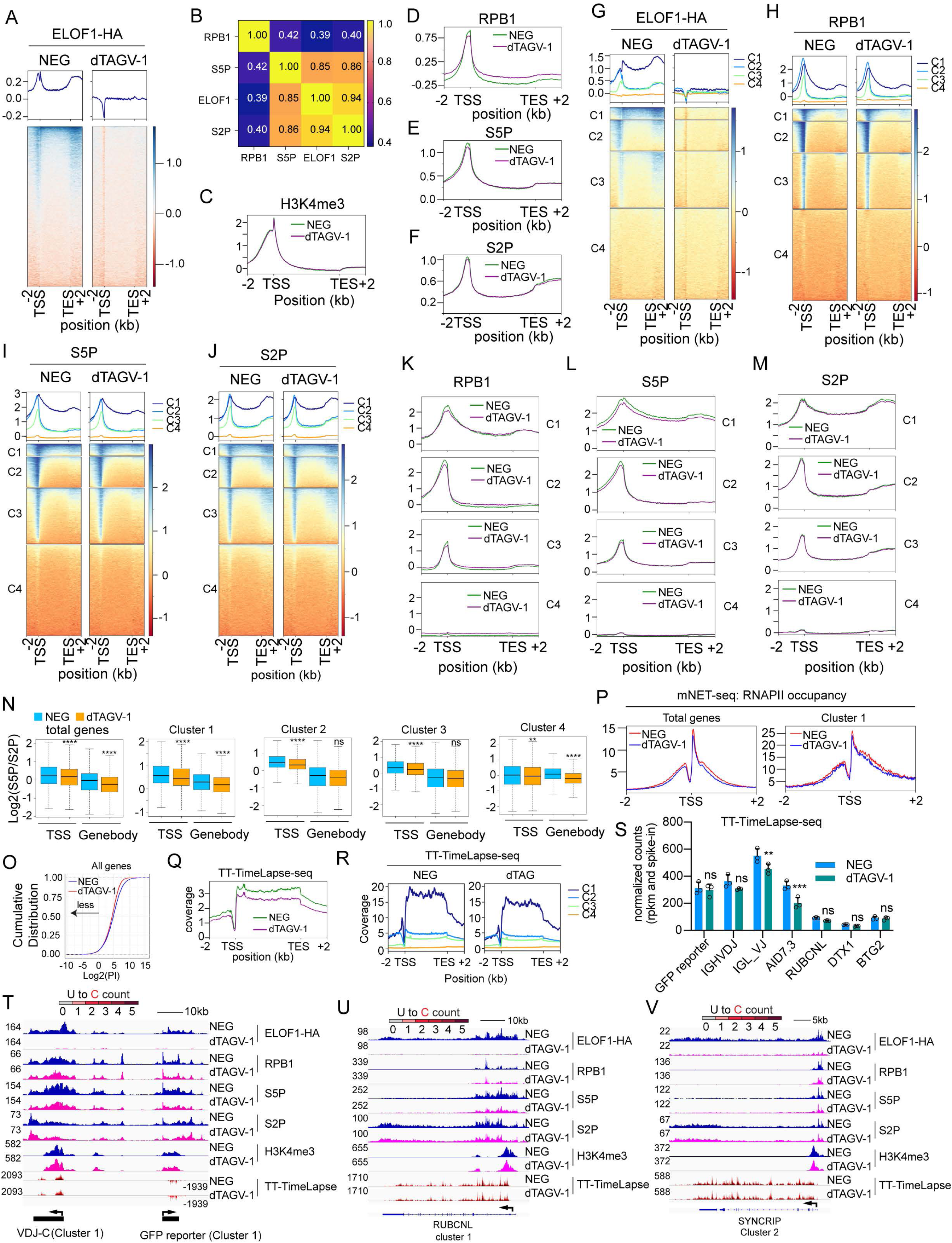
ELOF1 modulates RNAPII occupancy and pausing. (A) ELOF1 CUT&RUN-seq heatmap and profile were generated from ELOF1-degron cells treated with NEG or dTAGV-1 for 4 hours followed by 2-day dox treatment. TSS, transcription start site. TES, transcription end site. (B) Correlation plot using ELOF1, RPB1, S5P-RNAPII and S2P-RNAPII CUT&RUN-seq peaks. (C-F) CUT&RUN-seq metagene plot of H3K4me3 (C), RPB1 (D), S5P-RNAPII (E), S2P-RNAPII (F) in ELOF1-degron RASH-1C cells treated with NEG or dTAGV-1. (G-J) Unsupervised K clustering was performed based on ELOF1 and RPB1 CUT&RUN-seq using deepTools (K=4). Heatmap and profile for four clusters were generated for ELOF1 (G), RPB1 (H), S5P-RNAPII (I) and S2P-RNAPII (J). (K-M) Metagene plot of RPB1 (K), S5P-RNAPII (L), S2P-RNAPII (M) in four clusters. (N) Log2(S5P-RNAPII/S2P-RNAPII) in TSS or gene body in four clusters. Tukey’s HSD test was used to compare group means after finding significant differences with ANOVA. ****p<0.0001; ns, not significant. (O) RNAPII pausing index (PI) in ELOF1-degron cells treated with NEG or dTAGV-1 for 4 hours followed by 2-day dox treatment. Cumulative index plots of pausing index were calculated from RNAPII CUT&RUN-seq data. (P) Metagene profiles aligned at the TSS showing mean occupancy of RNAPII (mNET-seq) upon ELOF1 degradation. Sense profiles are shown. (Q) Metagene profile of TT-TimeLapse-seq in ELOF1-degron cells treated with NEG or dTAGV-1 for 4 hours followed by 2-day dox treatment. (R) Metagene profile of TT-TimeLapse-seq in four clusters. (S) Normalized counts of indicated genes by TT-TimeLapse-seq. Statistical significance was calculated using two-way ANOVA with Tukey’s multiple comparisons test. ***p value<0.001; **p value<0.01; ns, not significant. (T-V) Genome browser tracks of ELOF1, RPB1, S5P-RNAPII, S2P-RNAPII, H3K4me3 CUT&RUN-seq and TT-TimeLapse-seq in *IGH VDJ* (S), *GFP reporter* (S), *RUBCNL*(T) and *SYNCRIP* (U) regions. Reads for TT-TimeLapse-seq categorized by U to C mutations.

Unsupervised K means clustering was used to segregate genes into four clusters (C1-C4) based on ELOF1 and RPB1 CUT&RUN-seq data. Cluster 1 (1,189 genes; 6.3%) exhibited the highest overall levels of ELOF1 and RNAPII, cluster 2 had the highest levels of these factors at the TSS but much lower levels in the gene body, and clusters 3 and 4 showed progressively lower levels throughout the gene (Figures 4G and 4H). Similar trends were observed for S5P- and S2P-RNAPII (Figures 4I and 4J). Of the 45 protein coding genes previously identified as AID targets in Ramos,^37^ a majority (62%) are in cluster 1 (Figure S4D) as are *HTS7-GFP*, *IGH-VDJ, and IGL-VJ* (*RUBCNL*, *BTG2*, and *DTX1* are among the 45 previously identified loci). Cluster 1 is therefore highly enriched in AID target genes.

Comparison of ELOF1 and RNAPII profiles in cluster 1 reveals high levels of ELOF1 and all forms of RNAPII with only a modest peak at the TSS relative to the gene body, reduced S5P-RNAPII throughout the entire transcription unit upon degradation of ELOF1, and S2P-RNAPII profiles that are largely unchanged by loss of ELOF1 (Figures 4K-4M). While S2P is a mark of elongating RNAPII and is abundant in gene bodies, S5P is associated with paused RNAPII, tending to accumulate at the promoter proximal pause site immediately downstream of the TSS and declining in the gene body.^42, 43^ Loss of ELOF1 causes a drop in the ratio of S5P to S2P in the vicinity of the TSS and in the gene body that is particularly striking in cluster 1 (Figure 4N), suggesting a decrease in paused RNAPII. In support of this conclusion, loss of ELOF1 caused a substantial decrease in the RNAPII pausing index in clusters 1, 2 and 3 (Figures 4O and S4E). To investigate the change of RNAPII distribution in the absence of ELOF1 with higher resolution, we performed anti-RNAPII mNET-seq which detects the 3’ ends of Pol2-associated nascent transcripts and provides strand-specific mapping of chromatin-associated Pol2 at single nt resolution^44^. Consistently, we observed a small but detectable decrease in RNAPII occupancy in the promoter proximal region in all four clusters upon loss of ELOF1(Figure 4P and S4F), consistent with a role for ELOF1 in supporting RNAPII pausing. To determine the effects of ELOF1 on RNAPII RNA synthesis, we employed transient transcriptome sequencing with “timelapse chemistry” (TT-TimeLapse-seq)^45^ with a 5 min pulse of 4-thiouracil (4sU), using ELOF1 degron RASH-1 cells treated with NEG or dTAGV-1 and dox induced for 2 days. Metagene analysis showed that spike-in normalized TT-TimeLapse-seq coverage across the gene body was reduced ∼20% in cells treated with dTAGV-1 (Figure 4Q), with the reduction observed in all length groups of genes (0-50 kb, 50-100 kb, >100 kb) (Figure S4G). When combined with the observation that levels of S2P-RNAPII are not changed by loss of ELOF1, this reduction in RNA synthesis is in good agreement with two previous studies that found that RNAPII elongation rates are decreased ∼25% in *ELOF1* KO versus WT cells.^22, 23^ In genes over 50 kb long, we observed increasing divergence between the NEG and dTAGV-1 TT-TimeLapse-seq traces toward the 3’ ends of genes (Figure S4G), consistent with previous observations with *ELOF1* KO cells.^23^ This is likely explained by the cumulative effect of repeated perturbations in RNAPII elongation predicted to arise in the absence of ELOF1, for example upon encountering nucleosomes or natural pause sites.^22, 26^ Cluster 1 genes showed the highest levels of RNA synthesis, consistent with their high levels of S2P-RNAPII, and loss of ELOF1 reduced transcription comparably in all four clusters (Figures 4R and S4H). Notably, while SHM target genes showed somewhat reduced TT-TimeLapse-seq signals with dTAGV-1 treatment, the decrease was not statistically significant relative to NEG treatment for the majority of them (Figure 4S), arguing that reduced transcription levels per se do not explain the large decrease in SHM observed upon loss of ELOF1.

Genome browser tracks for cluster 1 SHM target genes illustrate the findings described above, including the lack of a strong peak of RNAPII near the TSS, broad peaks of all forms RNAPII that extend into the gene body, and reduced S5P-RNAPII and TT-TimeLapse-seq signals after ELOF1 degradation (Figures 4T, 4U and S4I, S4J). In contrast, a typical cluster 2 gene displays clear peaks of RPB1 and S5P-RNAPII at the TSS, coincident with the peak of H3K4me3, and much lower total and S5P-RNAPII in the gene body compared to cluster 1 (Figure 4V). We conclude that most Ramos SHM targets fall into a class of genes with high expression and low pausing index and that loss of ELOF1 further reduces pausing without affecting S2P-RNAPII levels.

## Discussion

RASH cell lines provide a useful tool for factor discovery and mechanistic studies related to SHM. The rapidity and ease with which SHM can be assayed is complemented by the high efficiency with which the Ramos genome can be manipulated. Ramos has been used extensively for the study of SHM for nearly three decades.^46, 47^ ELOF1 represents an experimentally tractable system for studying the link between RNAPII transcription and AID targeting because ELOF1 deficient cells are viable and proliferate well for extended periods. This stands in contrast to SPT5, degradation of which leads to a rapid, global destabilization of RNAPII and cell lethality,^38, 39^ making it difficult to disentangle general effects on RNAPII function from specific roles in SHM and CSR.

Our findings demonstrate that loss of ELOF1 strongly compromises AID-mediated deamination, SHM, and CSR, and that this is accompanied by a decrease in parameters of RNAPII pausing/stalling and in RNAPII RNA synthesis without a decrease in levels of S2P-RNAPII, likely due to reduced RNAPII elongation rates.^22, 23^ ELOF1 function in SHM and CSR requires RNAPII binding and AID can be recruited into complexes containing ELOF1 and RNAPII. Based on these observations, we consider a model for the mechanism of action of ELOF1 in SHM and CSR in which ELOF1 is important for the formation and/or stability of stalled RNAPII complexes that have been proposed to serve as the substrate for AID action (Figure S5). RNAPII stalling due to impediments such as transcriptional supercoiling, R-loops, AID generated template strand deoxyuridines, and RNAPII collisions, has been suggested to be an important step in SHM, CSR, and AID off-target activity^4, 12, 13, 35, 48–50^, and has been linked to the activity of SHM enhancer elements, also referred to as Diversification Activators (DIVACs).^15^ It has been proposed that the stalled RNAPII complex is subsequently destabilized by ubiquitin ligases and RNA exosome to allow AID access to both DNA strands.^5, 51, 52^ In the absence of ELOF1, the RNAPII pausing index is reduced and S5P-RNAPII is uniformly decreased at the TSS and gene body; however, the levels and distribution of H3K4me3 are unchanged, in keeping with the previous finding that transcription initiation is not affected by loss of ELOF1.^23^ Nor does loss of ELOF1 appear to effect release of elongation competent RNAPII into the gene body, as indicated by the unaltered levels of S2P-RNAPII. By selectively sustaining S5P-RNAPII complexes, ELOF1 could support the formation of an optimal RNAPII substrate for AID.

Our data are consistent with the possibility that ELOF1 also contributes to SHM and CSR by influencing the AID-RNAPII interaction. During TC-NER, ELOF1 makes direct contact with two TC-NER components surrounding the DNA entry channel to orchestrate conformational changes and a network of interactions needed for DNA repair.^24^ By analogy, ELOF1 might create, directly or indirectly, a platform for positioning of AID adjacent to the DNA entry channel.

### Limitations of the study

The contribution of ELOF1 to SHM and CSR has been assessed in Ramos and CH12F3 cells, respectively, which confirms its role in AID-mediated reactions in both human and mouse contexts but not in primary B cells. There is no evidence linking the TC-NER pathway to SHM or CSR and essential TC-NER factor CSB (Cockayne syndrome B) is not required for CSR.^53^ We cannot, however, rule out the possibility that ELOF1 acts through the TC-NER pathway specifically to support SHM. In ELOF1 degron experiments, it is possible that small amounts of ELOF1 persist in chromatin-bound RNAPII complexes, potentially leading us to underestimate the contribution of ELOF1 to SHM.

## Materials and Methods

### Cell culture

Ramos cells and cell lines derived from Ramos cells were grown in RPMI-1640 (Gibco 11875) supplemented with 10% FBS (Gemini), 0.5 mg/mL penicillin–streptomycin–glutamine (Gibco10378016) at 37°C, 5% CO_2_. 293T cells were grown at 37°C, 5% CO_2_ in DMEM (Gibco 10566) supplemented in the same fashion to Ramos cells. CH12 cells were cultured in RPMI 1640 media supplemented with 10% FBS, 1x MEM non-essential amino acids, 10 mM HEPES, 2 mM Glutamax, 1 mM sodium pyruvate, 55 μM 2-mercaptoethanol and penicillin and streptomycin.^54^

### Lentiviral and retroviral transduction

293T cells in 6-well plates were grown to 50–80% confluence and then transduced with 1 μg lentiviral plasmid, 0.6 μg psPAX2 (Addgene #12260), and 0.4 μg pMD2.G (Addgene #12259) or 1 ug retroviral plasmid and 1 ug packaging plasmid pkat2 with JetPrime reagent (Polyplus 114-07) according to the manufacturer’s protocol. At 48 hours after transfection, lentivirus or retroviral-containing media were collected and filtered through a 0.45 μm filter before being used to infect cells. Ramos cells were transduced as reported previously.^55^

### Genome-wide CRISPR screen

The LentiCRISPR-V2 pooled library (GeCKO v2) was amplified as described previously.^56^ A total of 200 million RASH-1 cells were transduced with a GeCKO v2.0 single-guide RNA (sgRNA) library containing 122,441 sgRNAs targeting 19,050 genes followed by puromycin selection (2 μg/ml) for 7 days. Then cells were treated with 200 ng/ml dox to induce AID7.3 expression for 4 days. Cells were sorted into two groups: GFP negative or GFP positive. Cellular DNA from GFP negative and GFP positive and unsorted cells was extracted using the DNeasy Blood & Tissue Kits (Qiagen, 69504) according to the manufacturer’s instruction. sgRNA sequences were amplified using High-Fidelity PCR master mix (NEB) and amplicons were purified from 2.5% agarose gel. Amplicons were sequenced using an Illumina HiSeq2500 (40 million reads per sample). The enrichment of genes in GFP negative versus GFP positive was ranked by the MAGeCK algorithm. ^57^ The screen was performed in triplicate.

### Generation of degron cell lines and AID off-targeting cell lines

ELOF1-degron cell lines were generated based on RASH-1C that constitutively expresses Cas9^20^ or WT Ramos cells. Plasmid expressing sgRNA targeting *ELOF1 stop codon* region and donor plasmid containing left and right homology arms attached to HA-FKBP12^F36V^-mScarlet sequences^58^ were delivered into RASH-1C cells or WT Ramos cells by electroporation using Amaxa Nucleofector II (Lonza) with program O-006 according to the manufacturer’s instructions. Homemade 1M buffer was used for electroporation according to.^59^ 5 days later, mScarlet positive cells were sorted into a 96 well plate to obtain single cell clones. Expanded clones were cultured for 15 days in a 96 well plate and then analyzed by FACS to detect mScarlet signal and then validated by Sanger sequencing.

AID off-targeting cell lines were generated according to the method described in^20, 36^. 1 million AID7.3in^55^ cells were infected with virus expressing *GFP7-E* vector at MOI < 0.01. Two days post infection, blasticidin with final concentration of 5 μg/ml was added. Two weeks later, GFP-positive cells were sorted in a single-cell sort mode into 96-well tissue culture plates to obtain single cell clones. Expanded clones were cultured for 18 days in 96 well plates and then 40 µL cells were transferred to a new plate and then treated with 200 ng/ml dox for 6 days, followed by GFP loss analysis. The clones that yielded a high percentage of GFP loss by dox treatment were selected. The integration site of *GFP7-E* was identified using the splinkerette-PCR method described in^36^.

### IgM and GFP loss assay

For the IgM loss assay in RASH-1 cells, ∼1.0 × 10^6^ cells treated with or without 200 ng/ml dox were stained with APC Mouse Anti-Human IgM (BD 551062) diluted 1:100 in FACS Buffer (1× PBS with 2% FBS) and incubated for 20 min at RT. For the IgM loss assay in WT Ramos, cells were initially sorted to be IgM^+^, cultured for 5 days and the percentage of IgM^−^ cells was measured by flow cytometry. For the GFP loss assay, RASH-1C cells, RASH-1C derived cells, and AID off-targeting reporter cells treated with or without 200 ng/ml dox for 2-5 days as indicated in the figures were assayed for the percentage of GFP^+^ and GFP^−^ cells by flow cytometry.

### High throughput sequencing

Analyses of mutations, deletions, and insertions were performed as described in^20^. Briefly, Genomic DNA was extracted using DNeasy Blood & Tissue Kit (Qiagen, 69504) according to the manufacturer’s instructions. *HTS7* was amplified using the oligos: (forward) TCGTCGGCAGCGTCAGATGTGTATAAGAGACAGTTCCAAAGTAGACCCAGCCTTCTAA and (reverse) GTCTCGTGGGCTCGGAGATGTGTATAAGAGACAGGATTCTCCTCCACATCACCACAG. *IGH VDJ* was amplified using the oligos: (forward) TCGTCGGCAGCGTCAGATGTGTATAAGAGACAGAGGAATGCGGATATGAAGATATGAG and (reverse) GTCTCGTGGGCTCGGAGATGTGTATAAGAGACAGAGTAGCAGAGAACAGAGGCCCTAGA. *IGLV2-14J2* was amplified using the oligos: (forward) TCGTCGGCAGCGTCAGATGTGTATAAGAGACAGCACTGACTCACTGGCATGTATTTCT and (reverse) GTCTCGTGGGCTCGGAGATGTGTATAAGAGACAGGCTGACCACAAGTTGAGACAAGATA. *AID7.3* was amplified using the oligos: (forward) TCGTCGGCAGCGTCAGATGTGTATAAGAGACAG TGGAGCAATTCCACAACACT (reverse) GTCTCGTGGGCTCGGAGATGTGTATAAGAGACAG GAATTTTCATGCAGCCCTTC *RUBCNL* was amplified using the oligos: (forward) TCGTCGGCAGCGTCAGATGTGTATAAGAGACAG GATGGGGGTAGGGTGAAGAC (reverse) GTCTCGTGGGCTCGGAGATGTGTATAAGAGACAG AAATCATTTTATTCGTCCCATGA *DTX1* was amplified using the oligos: (forward) TCGTCGGCAGCGTCAGATGTGTATAAGAGACAG GAACTGGGTACTGGCAGGAG (reverse) GTCTCGTGGGCTCGGAGATGTGTATAAGAGACAG GAGGAAGGGGAGGAGACAGA *BTG2* was amplified using the oligos: (forward) TCGTCGGCAGCGTCAGATGTGTATAAGAGACAG CTCCTTTCAGAGCTCTCAGTCC (reverse) GTCTCGTGGGCTCGGAGATGTGTATAAGAGACAG CTCACCTGTGAGTGCCTCCT *BCL6* was amplified using the oligos: (forward) TCGTCGGCAGCGTCAGATGTGTATAAGAGACAG GACAGCCGCTTTGGATAAC (reverse) GTCTCGTGGGCTCGGAGATGTGTATAAGAGACAG GAGAAACGCGCCTCTGTTC *MYC* was amplified using the oligos: (forward) TCGTCGGCAGCGTCAGATGTGTATAAGAGACAG CAGTGCGTTCTCGGTGTG (reverse) GTCTCGTGGGCTCGGAGATGTGTATAAGAGACAG TTTTATACTCAGCGCGATCC *BACH2* was amplified using the oligos: (forward) TCGTCGGCAGCGTCAGATGTGTATAAGAGACAGTGGCAACACAAAGCCAGTAG (reverse) GTCTCGTGGGCTCGGAGATGTGTATAAGAGACAGTTTCAGTGTCTCTGGTGTGGA *BCL7A* was amplified using the oligos: (forward) TCGTCGGCAGCGTCAGATGTGTATAAGAGACAG GAGGTCACCCCAGACTAGCA (reverse) GTCTCGTGGGCTCGGAGATGTGTATAAGAGACAG CTGCAAACTGGACCCTCAGT

PCR reactions were performed using NEBNext^®^ High-Fidelity 2X PCR Master Mix (NEB) under the conditions: 98°C for 3 min, followed by 25 cycles at 98°C for 10s, 60°C for 30s, 72°C for 30s, and 72°C for 2 min in a 25 µL reaction. PCR products were multiplexed using nextera XT index kit (Illumina) and applied to an Illumina MiSeq platform using MiSeq reagent kit V3 or NextSeq 2000 platform using NextSeq1000/2000 P1 Reagents (600cycles, 20075294, Illumina) (performed by Yale Center for Genome Analysis).

### Proximity ligation assay (PLA)

PLA was performed according to the manufacturer’s protocol (Sigma, DUO92101, DUO92013, DUO94004) with minor modifications. Briefly, 1 million cells were harvested and resuspended in nuclei extraction buffer (0.32 M sucrose, 3 mM CaCl_2,_ 2 mM magnesium acetate, 0.1 mM EDTA, 10 mM Tris-HCl, pH 8.0, 1 mM DTT, 0.5 mM PMSF, 0.5% IGEPAL). Nuclei were fixed, permeabilized, blocked, and incubated with primary antibodies overnight. The Duolink PLA probe incubation, ligation, and amplification were performed before detection of PLA signals by using Duolink^®^ In Situ Detection Reagents FarRed kit (imaging) or by Duolink^®^ flowPLA Detection Kit -FarRed (FACS). Images were acquired using a Leica SP8 laser scanning confocal microscope (West Campus Imaging Core, Yale). The number of PLA foci per cell was quantified by Imaris.

### Co-immunoprecipitation (Co-IP)

Co-IP was performed as previous described with modifications.^8^ 40 million cells were resuspend with 1 ml nuclear extraction buffer (20 mM HEPES pH 7.9, 10 mM KCl, 0.1% Triton X-100, 20% glycerol, 1 x protease inhibitor cocktail (Roche)) and rotated at 4°C for 20 min. Nuclear pellets were then resuspended in 400 μL of low salt buffer (20 mM HEPES pH 7.5, 10 mM KCl, 1 mM MgCl2, 10% glycerol, 1% NP40, 1 x protease inhibitor cocktail (Roche)) containing 250 U/ml Benzonase nuclease (Sigma) and were sonicated 5 x 10 s at 50% amplitude at 4°C. Samples were then incubated 30 min on a rotator at 4°C followed by centrifuged at max speed for 10 min. Supernatant was collected and mixed with 250 μL of high salt buffer (low salt buffer containing 400 mM NaCl). 2.5 μg anti-AID antibody (Invitrogen, 39-2500) was added and samples were incubated overnight. Protein complexes were pulled down by 4 h incubation with 25 μL Pierce™ Protein A/G Magnetic Beads (ThermoFisher, 88802) and the samples were prepared by boiling in Laemmli–SDS sample buffer for subsequent analysis by western blotting.

### Class switch recombination

For CSR from IgM to IgA in the CH12F3 cell line, cells were plated at a density of 1×10^5^ per mL of media and stimulated with anti-CD40 (1 μg/mL, clone 1C10, Biolegend), rmIL-4 (10 ng/mL, Peprotech), and rhTGF-β (1 ng/mL, R&D) (CIT) for 72 hours.

### CUT&RUN-seq

CUT&RUN was performed according to the manufacturer’s protocol using CUTANA™ ChIC/CUT&RUN Kit (Epicypher, 14-1048). Briefly, ELOF1-degron cells were treated with 0.5 μM NEG (Tocris, 6915) or dTAGV-1 (Tocris, 6914) for 4 hours followed by 200 ng/ml dox treatment. After 2 days, 7.5×10^5^ cells were harvested for nuclei extraction. Isolated nuclei were then bound to activated concanavalin A-coated magnetic beads and then incubated with 0.5 μg HA antibody (Santa Cruz, sc-7392), Rpb1 CTD (Cell Signaling Technology, 2629, 1:50), S2P-RNAPII (Cell Signaling Technology, 13499, 1:50) or S5P-RNAPII (Cell Signaling Technology, 13523, 1:50) or 0.5 μg H3K4me3 (Epicypher) at 4°C overnight on a nutator. After two washes with cell permeabilization buffer, 2.5 μl pAG/MNase was added to beads resuspended in 50 μl cell permeabilization buffer and incubated for 10 min at room temperature. After two washes with cell permeabilization buffer, 1 μl 100 mM CaCl_2_ was added and then incubated at 4°C for 2 hours on a nutator. 33 μl stop buffer was added and incubated at 37°C for 10 min, and DNA was purified using SPRIselect reagent (beads). Libraries were prepared using the NEBNext Ultra II DNA Library Prep Kit (NEB, E7645S) and sequenced on an Illumina NovaSeq X (paired-end, 2 x 75 bp) at an average depth of 20 million reads per sample.

### mNET-seq

mNET-seq was performed as previously described with minor modifications.^60^ 25 million cells were washed with ice-cold DPBS once, resuspended in 4 ml of ice-cold HLB + N buffer (10 mM Tris-HCl (pH 7.5), 10 mM NaCl, 2.5 mM MgCl2, 0.5%(v/v) NP-40 and proteinase inhibitor) and incubated on ice for 5 min. The cell suspension was then underlaid with 1 ml of HLB + NS buffer (10 mM Tris-HCl pH 7.5, 10 mM NaCl, 2.5 mM MgCl2, 0.5% (v/v) NP-40, 10% (w/v) sucrose and 1* proteinase inhibitor) and centrifuged to pellet the nuclei at 400g, 5 min at 4°C. The nuclei were resuspended in 125 μl of NUN1 lysis buffer (20 mM Tris-HCl pH 8.0, 75 mM NaCl, 0.5 mM EDTA, 50% (v/v) glycerol and 1* proteinase inhibitor), transferred to a 1 ml tube and to each tube was added 1.2 ml NUN2 buffer (20 mM HEPES-KOH pH 7.6, 300 mM NaCl, 0.2 mM EDTA, 7.5 mM MgCl_2_, 1% (v/v) NP-40, 1 M urea and 1* proteinase inhibitor) to precipitate the chromatin, and the sample was incubated on ice for 15 min with occasional vortexing. The lysates were then centrifuged at 16,000g for 10 min at 4°C to pellet the chromatin and the chromatin pellets were washed with 100 μl 1* MNase buffer and then digested with 2 μl MNase (CST) for 2 min at 37°C with mixing at 1,400 rpm on a thermomixer. The digestion was stopped by adding 10 μl of 500 mM EDTA and transferred onto ice for 10 min. Reactions were centrifuged at 16,000g for 5 min at 4°C and the supernatant was diluted with 1 ml of NET-2 buffer (50 mM Tris-HCl pH 7.4, 150 mM NaCl, 0.05% (v/v) NP-40). 1 ml beads and antibody (5 μg antibody conjugated to 75 μl protein A/G beads) were added and the mixture was incubated in the cold room for 1.5 h. This was followed by eight washes with NET-2 buffer and one wash with 500 μl of PNKT buffer containing 1* T4 polynucleotide kinase (PNK) buffer (NEB, M0201L) and 0.1% (v/v) Tween-20. The beads were incubated in 100 μl of PNK reaction mix containing 1* T4 DNA ligase buffer (containing ATP), 0.1% (v/v) Tween- 20, and T4 PNK 3′ phosphatase minus (NEB, M0236L) at 37°C for 10 min. Beads were washed with 1 ml of ice-cold NET-2 buffer by inverting the tube, immersed in Trizol, and 1 ng of Drosophila mRNA spike-in was added to each sample. The RNA was extracted in Trizol (200 μl) and libraries were generated with QIA miRNA kit seq (Qiagen 331505). Libraries were sequenced on an Illumina NovaSeq (paired end 2*100) at an average depth of 50 million reads per sample.

### TT-TimeLapse-seq

TT-TimeLapse-seq was performed as previously described.^45^ Approximately 20 million cells per sample were labeled with 1 mM 4sU (Fisher Scientific, 13957-31-8) for 5 min. Cells were harvested and washed in ice-cold PBS and dissolved in Trizol. Total RNA was extracted and treated with TURBO™ DNase (ThermoFisher, AM2238) to deplete genomic DNA. 5% genomic DNA depleted Drosophila S2 cell spike-in RNA (5 min 4sU treated) was added. RNA was then sheared by mixing with 2x RNA fragmentation buffer (150 mM Tris pH 7.4, 225 mM KCl, 9 mM MgCl_2_) and heated to 94°C for 3.5 min. RNA shearing was stopped by adding EDTA (to a final concentration of 50 mM) and incubated on ice for 2 min. RNA was purified using RNeasy Mini Kit (Qiagen, 74104). Fragmented RNA was biotinylated using 10% MTSEA-biotin-XX (VWR, 89139-636) prepared in DMF. The biotinylated RNA was purified and isolated using 10 μl Dynabeads MyOne Streptavidin C1 magnetic beads (Invitrogen, 65001). 4sU-biotinylated RNA was eluted by reducing the disulfide bond that formed between biotin and 4- thiouridine, thereby eluting the s^4^U-RNA and leaving biotin bound to the streptavidin beads using 25 μl elution buffer (100 mM DTT, 20 mM DEPC, pH7.4, 1 mM EDTA, 100 mM NaCl, 0.05% Tween-20). TimeLapse chemistry was carried out as describe in^45^ to convert U to C. Final RNA was converted to libraries using the Takara Pico Mammalian V3 kit (Takara, 634938). Libraries were sequenced on an Illumina NovaSeq X at an average depth of 30 million reads per sample.

### User-S1-END-seq

USER-S1-END-seq protocol was adapted from the END-seq and S1-END-seq protocols previously described.^33, 61–63^ To compare samples accurately, a spike-in control was added to USER-S1-END-seq samples. The spike-in consists of a G1-arrested mouse Abelson transformed pre-B cell line (Lig4−/−) carrying a single zinc-finger-induced DSB at the TCRβ enhancer.33 Breaks at the TCRβ enhancer are present in all cells, which were mixed in at a 20% frequency with Ramos cells for data normalization. 30 million Ramos cells mixed with spike-in control were embedded in agarose plugs and treated with Proteinase K for 1h at 50°C and 7 hours at 37°C. After protein digestion, the agarose plugs were washed two times in washing buffer (10mM Tris-HCl, pH 8.0, 50mM EDTA) and then four times in TE (10mM Tris-HCl, pH 8.0, 1mM EDTA), which was followed by RNase A treatment for 1 hour at 37°C. The plugs were then washed with washing buffer once and then three times with EB buffer (10mM Tris-HCl, pH 8.0), pre-incubated with CutSmart buffer under agitation 400 rpm for 10 minutes at room temperature and then incubated with USER reaction, 5U of USER (NEB Cat#M5505S) diluted in CutSmart buffer, for 30 minutes at 37°C under agitation 400 rpm. The USER reaction was then removed, the plugs were then rinsed with wash buffer and then washed once for 10 minutes with wash buffer prior to three 10 minutes washes with EB buffer. This was followed by S1 nuclease treatment where the plugs were washed once with washing buffer for 10 minutes, washed twice with EB buffer for 10 minutes each and then equilibrated with two washes of 10 min with S1 nuclease buffer (40 mM sodium acetate pH 4.5, 300 mM NaCl, 2 mM ZnSO4). Plugs were then incubated at 37°C for 30 min with 2 U of S1 nuclease (Sigma Aldrich Cat#EN0321) in 100 μL per plug diluted in S1 nuclease buffer. Finally, the DNA ends were blunted with exonuclease VII (NEB) for 1 hour at 37°C and exonuclease T (NEB), which was followed by adapter ligation, DNA sonication, and library preparation performed as described previously.^33, 64^ Sequencing was performed on the Illumina NextSeq 2000 (100 bp single-end reads) or NextSeq 550 (75 bp single-end reads).

### Bioinformatic analyses: CRISPR screen analysis

We used MAGeCK^57^, a comprehensive CRISPR screen analysis pipeline widely employed to identify essential gene hits. Briefly, following deep sequencing, reads were adapter-trimmed and mapped to the original sgRNA library. Counts across groups were normalized for library sizes and count distributions. A mean-variance model estimated the variance of over-dispersed sgRNA abundance, and a negative binomial distribution, similar to the edgeR package, tested for significant differences between treatment and control groups. sgRNAs were ranked, and a robust ranking aggregation (RRA) method was used to identify positively and negatively enriched genes. False discovery rate (FDR) was calculated via permutation tests.

### Mutation-seq analysis

Sequence reads were processed using a custom pipeline. Paired-end reads were merged using the fastq-join tool, requiring at least a 10-bp overlap with a mismatch rate of ≤8%. Merged reads were aligned to their respective reference sequences using the Burrows-Wheeler Aligner (BWA) with default settings. Python utility Pysamstats was utilized to compute statistics against reference sequence positions based on the aligned BAM files. Position-wise statistics were generated for reads with a mapping quality of at least 60 and a minimum base quality of 30. For each sample, the aligned BAM file was converted to TSV format using the Java-based utility sam2tsv. In-house AWK script generated per-read summary files for each observed variation, including deletions, insertions, and point mutations. Only nucleotide bases with a Phred quality score of ≥30 were considered for insertions and point mutations. For deletions, only reads with an average Phred score of ≥30 were included, as the relevant bases were deleted. We then identified WRC sites (AID hotspots) from the position-based statistics files generated by Pysamstats using R script.

### USER-S1-END-seq genome alignment and data visualization

Raw reads were aligned to the human genome (hg19) using Bowtie (v1.3.1) with -n 3 -l 50 -k1. Then, Samtools (v1.5.1) functions “view” and “sort” were used to convert and sort the aligned .sam files to sorted bam files. The mapped reads from .bam files were converted to .bed files using the bamTobed command from bedtools (v2.3.0). For data visualization, .bed files were converted to BedGraph using bedtools genomecov. Finally, BedGraph files were transformed into BigWig files using BedGraphToBigWig. Genome browser profiles were normalized to the library size (reads per million - RPM). Visualization of genomic profiles was done using IGV software v2.9.2.

For spike-in normalization, reads from the samples were aligned to both human (hg19) and mouse (mm10) genomes. The function “genomecov” from bedtools was used to normalized read density (reads per million, RPM). The scaling factor for the spike-in normalization was calculated using the number of reads mapped to the spike-in site in the mouse genome and the total number of reads mapped to the human genome. To generate the spike-in normalized reads, the normalized read density (RPM) was divided by the scaling factor.

### CUT&RUN-seq analysis

Paired-end reads were aligned to a modified version of the human genome (hg19) using Bowtie2 with local, very-sensitive alignment settings to ensure high-quality mapping. All downstream analyses was performed using the properly mapped read pairs. We merged three replicates and employed deepTools (version 3.5.1) to generate RPKM-normalized, input-subtracted (IgG control) bigwig files. These bigwig files were subsequently used to create all coverage profiles and other metaplots.

### Calculation of pausing index from RNAPII CUT&RUN-seq data

To calculate the pausing index (PI) from RNAPII CUT&RUN-seq data, we adapted the method from ^65^. For each annotated RefSeq isoform, the input-subtracted RNAPII signal was determined by overlaying the filtered RNA POL II reads to the TSSR (–50 bp to +300 bp around TSS) and the gene body (+300 bp downstream of TSS to +3 kb past TES) and the pausing index (PI) was calculated as

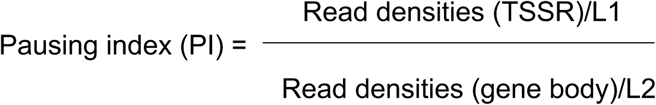

Read densities (TSSR)/L1 Read densities (gene body)/L2 where the TSSR length (L1) is always 350 bp and the gene body length (L2) is from +300 bp past TSS to 3 kb past TES. To consolidate PI values for genes with multiple RefSeq isoforms, we selected the isoform with the strongest RNAPII signal in the TSSR, provided it had at least 0.001 rpm/bp. For isoforms sharing the same TSS, we chose the longest isoform to capture the full RNAPII signal.

### mNET-Seq data analysis

The mNET-seq reads were processed by first extracting unique molecular identifiers (UMIs) using UMI-tools,^66^ followed by trimming of adapter sequences with Cutadapt. The reads were then aligned to a modified hg38 genome, merged with the Drosophila genome using the STAR aligner.^67^ Duplicates were removed with the UMI_Collapse tool.^68^ RNAPII positions were determined from the end of R2 reads for genes on the positive (+) strand and from the start of R1 reads for genes on the negative (-) strand. To account for coverage variability, counts from Drosophila genes were used to estimate scaling factors via DESeq2,^69^ which were applied to normalize the coverage profiles.

### TT-TimeLapse-seq analysis

TT-TimeLapse-seq reads from ELOF1 degron cells treated with NEG or dTAGV-1, containing spiked-in RNA from Drosophila cells were processed using the Snakemake implementation^70^ of the TT-TimeLapse pipeline as described in^45^. Briefly, the reads were trimmed to remove adaptor sequences using Cutadapt and were mapped to Bowtie2. Bam files containing uniquely mapped reads were, sorted and indexed using SAMtools. We used deepTools (V-3.5.1) to generate metaplots from the RPKM normalized and spike-in scaled bigwig files. The spike-in scaling factors were calculated using edgeR’s TMM strategy.^71^

To assess the differential RNA synthesis between the NEG and dTAGV-1 treated samples, the cutadapt trimmed reads were mapped to a combined hg19 and dm6 genome using HISAT-3N134 with default parameters and U-to-C mutation calls. Normalized mutation-specific coverage tracks were generated and were visualized using IGV.^72^ Drop-off probability for each gene is calculated as described in.^73^

### Statistical analysis

Data were subjected to statistical analysis and plotted using Graphpad Prism. Single comparisons were performed using the two-tailed Student’s *t* test, whereas multiple comparisons were assessed by one-way ANOVA with the Dunnett’s multiple comparison test or two-way ANOVA with Tukey’s multiple comparisons test. For all analyses, * p-value <0.05, ** p-value <0.01, ***p-value <0.001, ****p-value <0.001. Results are reported as mean ± SD as indicated in the figure legends.

## Data availability

The sequencing data sets generated during this study will be made available.

## Acknowledgements

We thank Yale Flow Cytometry Core for their assistance with cell sorting and analysis services. The Core is supported in part by an NCI Cancer Center Support Grant NIH P30 CA016359. DNA sequencing was performed by the Yale Center for Genome Analysis, which is supported by the National Institute of General Medical Sciences of the National Institutes of Health under Award Number 1S10OD030363-01A1. We thank the Yale West Campus Imaging Core for support and assistance in this work. Model diagram of Figure S5 was created with BioRender (https://biorender.com/) under the academic subscription of Yale School of Medicine, Department of Immunobiology. A.N. and G. M-R. are supported by the Intramural Research Program of the NIH, funded in part with federal funds from the National Cancer Institute under contract HHSN2612015000031. W.W. is supported by National Natural Science Foundation of China 32370575 and Chinese Academy of Sciences (318GJHZ2023004MI). This work was funded by research funds from NIH Shared Equipment grant #1S10OD028669-01 and NIH Grants R01-AI127642 (D.G.S.) and R01-GM137117 (M.D.S).

## Author Contributions

L.W. and D.G.S. designed most of the experiments, which were performed by L.W. Most computational analyses were performed by A.D.Y. G.M-R., W.W., and A.N. designed the USER-S1-END-seq experiments, which were performed by G.M-R., with computational analysis by W.W. D.X. provided the CRISPR loss of function sgRNA library in amplified form. A.P.P-U and M.D.S. provided Drosophila S2 cell spike-in RNA and assisted in analysis and interpretation of TT-TimeLapse-seq data. L.W. assembled the figures and L.W. and D.G.S. wrote the paper with input from other authors.

## Figure legends

**Figure S1.**
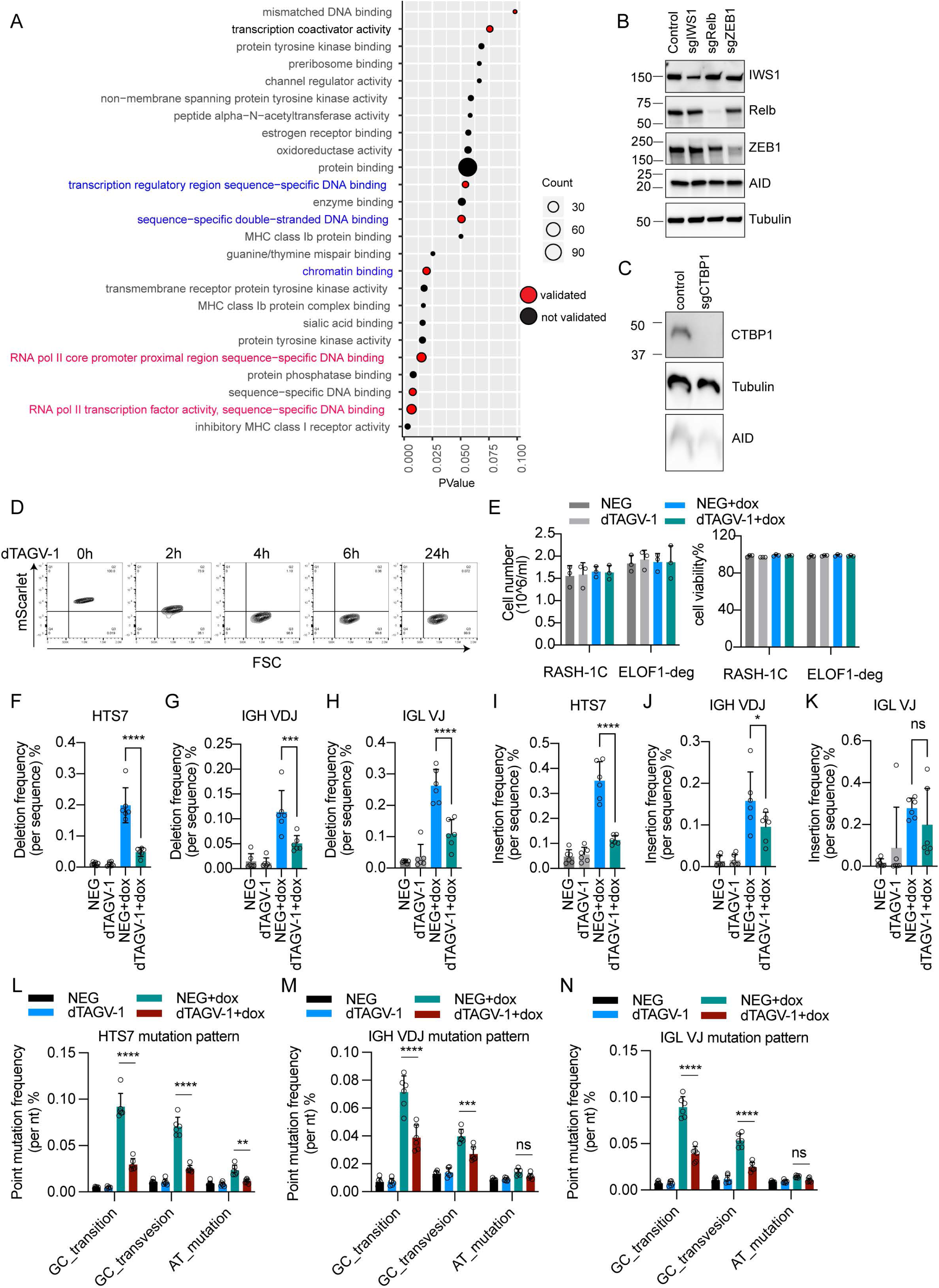
Related to Figure 1. (A) Gene ontology analysis of top 200 CRISPR screen hits ranked by MAGeCK. (B) Western blot showing protein levels of IWS1, Relb, ZEB1 and AID in RASH-1C cells treated with indicated sgRNAs. (C) Western blot showing CTBP1 protein in RASH-1C cells treated with CTBP1 sgRNA. (D) mScarlet signal in ELOF1-degron RASH-1C cells with indicated time-course of dTAGV-1 treatment. (E) Cell number and cell viability in WT RASH-1C cells and ELOF1-degron RASH-1C cells treated with dTAGV-1 for 2 days. (F-H) Deletion frequencies in *HTS7* (F), *IGH VDJ* (G) and *IGL VJ* (H) regions from ELOF1-degron RASH-1C cells treated with dTAGV-1 for 4 hours followed by 2-day dox or no dox treatment. (I-K) Insertion frequencies in *HTS7* (I), *IGH VDJ* (J) and *IGL VJ* (K) regions from ELOF1-degron RASH-1C cells treated with dTAGV-1 for 4 hours followed by 2-day dox or no dox treatment. (L-N) Mutation frequencies for G/C transitions, G/C transversions, and A/T mutations in *HTS7* (L), *IGH VDJ* (M) and IGL VJ (N) regions from ELOF1-degron RASH-1C cells treated with dTAGV-1 for 4 hours followed by 2-day dox or no dox treatment. Throughout the figure, data are presented with bar representing mean and error bar as ± SD. Statistical significance was calculated using one way ANOVA with Dunnett’s post-test for F-K and Two-way ANOVA with Tukey’s multiple comparisons test for L-N. ****p value<0.0001, ***p value<0.001; **p value<0.01; *p value<0.05; ns, not significant.

**Figure S2.**
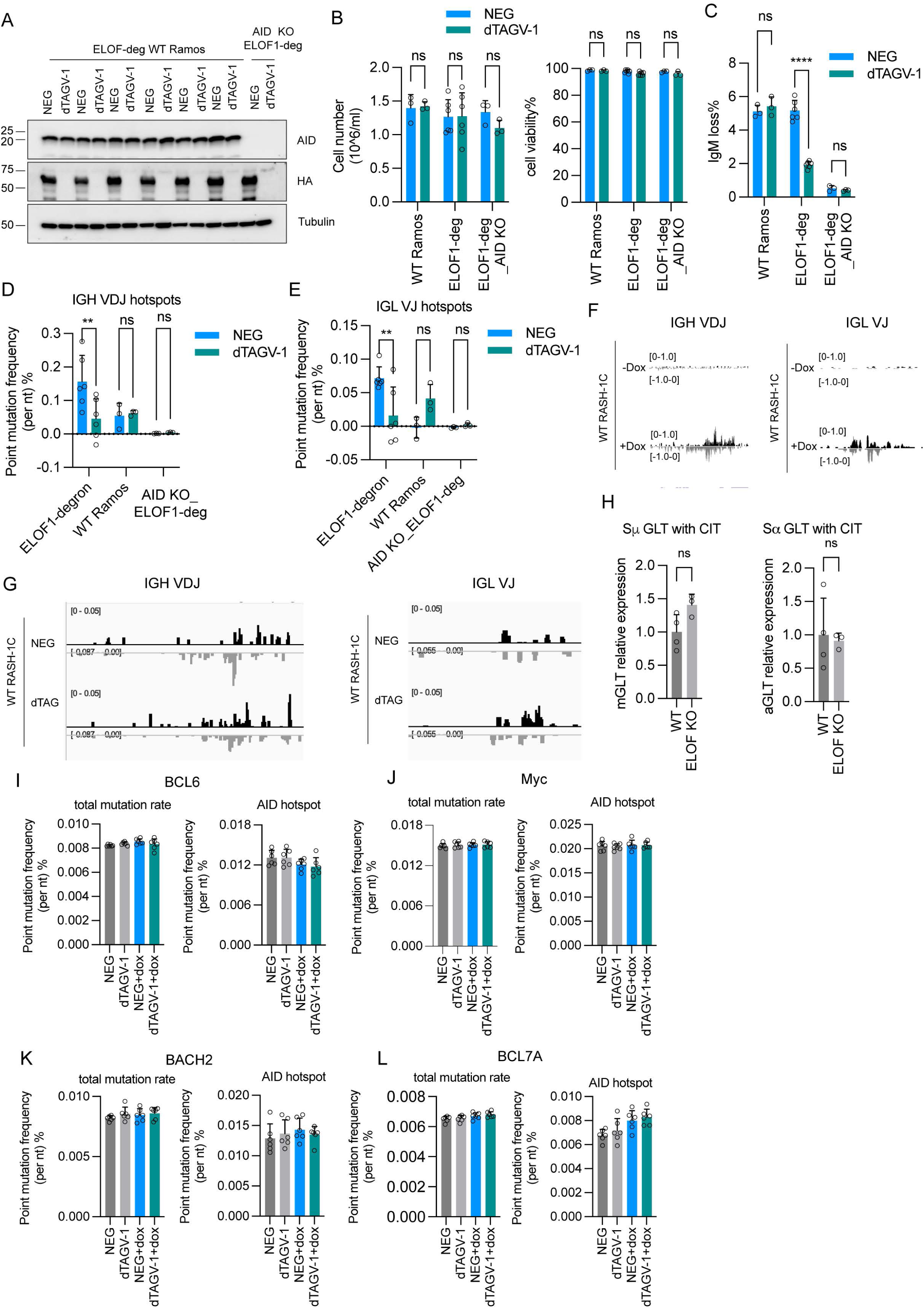
Related to Figure 2. (A) Western blot of six independent ELOF1-degron WT Ramos clones and *AID* KO ELOF1-degron WT Ramos treated with dTAGV-1 for 5 days. (B) Cell number and cell viability in ELOF1-degron WT Ramos cells treated with dTAGV-1 for 5 days. (C-E) WT Ramos cells, ELOF1-degron WT Ramos and *AID* KO ELOF1-degron WT Ramos were sorted for IgM positive cells. One day later, sorted cells were treated with NEG or dTAGV-1. After 5 days, cells were harvested for IgM loss (C) and mutation-seq for *IGH VDJ* (D) and *IGL VJ* regions (E). Cells before dTAGV-1 treatment (day 1) were also sequenced and provided a measure of the background mutation frequency. Point mutation frequency was calculated as point mutation frequency at day 5 minus point mutation frequency at day 1. (F) USER-S1-END-seq tracks showing peaks in *IGH VDJ* and *IGL VJ* regions from WT RASH-1C cells with or without 2-day dox treatment. (G) USER-S1-END-seq tracks showing peaks in *IGH VDJ* and *IGL VJ* regions from WT RASH-1C cells with or without dTAGV-1 treatment. (H) qPCR quantifying relative expression of GLT (germline transcription) in Sμ and Sα in cells treated with CIT for 3 days. (I-L) Total point mutation frequency and AID hotspot point mutation frequency in *BCL6* (I), *Myc* (J), *BACH2* (K) and *BCL7A* (L) regions from ELOF1-degron RASH-1C cells treated with dTAGV-1 for 4 hours followed by 5-day dox or no dox treatment. Throughout the figure, data are presented with bar representing mean and error bar as ± SD. Statistical significance was calculated using one way ANOVA with Dunnett’s post-test for I-L and Two-way ANOVA with Tukey’s multiple comparisons test for B-E and Student’s t-test for H. ****p value<0.0001; ***p value<0.001; **p value<0.01; ns, not significant.

**Figure S3.**
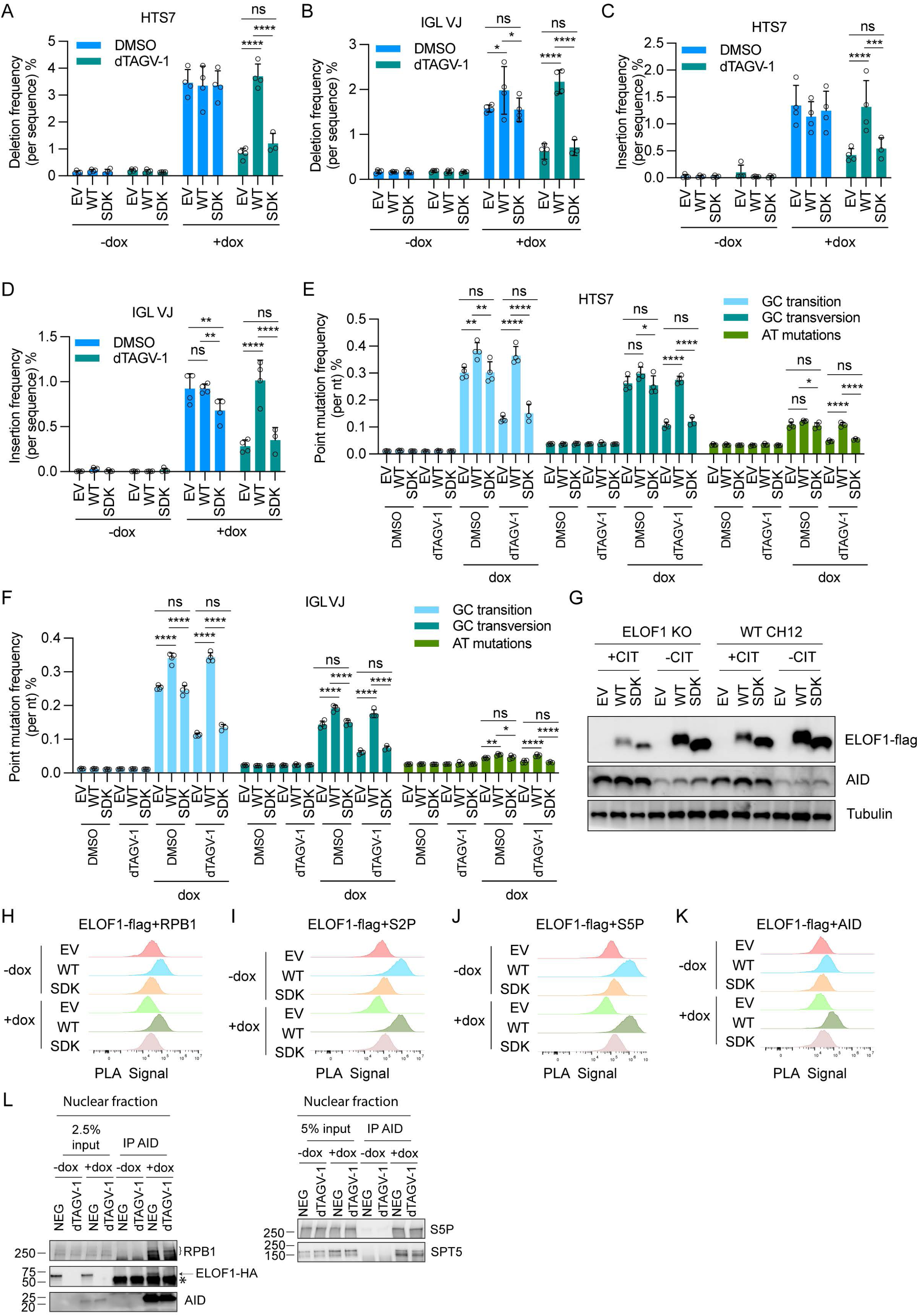
Related to Figure 3. (A, B) Deletion frequencies in *HTS7* (A), and *IGL VJ* (B) regions from ELOF1-degron RASH-1C cells reconstituted with WT ELOF1 or SDK mutant treated with dTAGV-1 for 4 hours followed by 4-day dox or no dox treatment. (C, D) Insertion frequencies in *HTS7* (C), and *IGL VJ* (D) region from ELOF1-degron RASH-1C cells reconstituted with WT ELOF1 or SDK mutant treated with dTAGV-1 for 4 hours followed by 4-day dox or no dox treatment. (E, F) Mutation frequencies for G/C transitions, G/C transversions, and A/T mutations in *HTS7* (E) and *IGL VJ* (F) regions from ELOF1-degron RASH-1C cells reconstituted with WT ELOF1 or SDK mutant treated with dTAGV-1 for 4 hours followed by 4-day dox or no dox treatment. (G) Western blot showing ELOF1 and AID protein levels in WT and *ELOF1* KO CH12F3 cells reconstituted with WT ELOF1 or SDK mutant and induced with CIT for 3 days. (H-K) PLA signal quantified by FACS in ELOF1-degron RASH-1C cells reconstituted with WT ELOF1 or SDK mutant treated with dTAGV-1 for 4 hours followed by 1-day dox or no dox treatment. (L) Anti-AID pull down from ELOF1-degron RASH-1C cells treated with NEG or dTAGV-1 for 4 hours followed by 1-day dox or no dox treatment. The arrow points to ELOF1 protein. Asterisk, background signal likely arising from the Ig heavy chain of the antibody used for pull down. n=1. Throughout the figure, data are presented with bar representing mean and error bar as ± SD. Statistical significance was calculated using Two-way ANOVA with Tukey’s multiple comparisons test for A-F. ****p value<0.0001; ***p value<0.001; **p value<0.01; ns, not significant.

**Figure S4.**
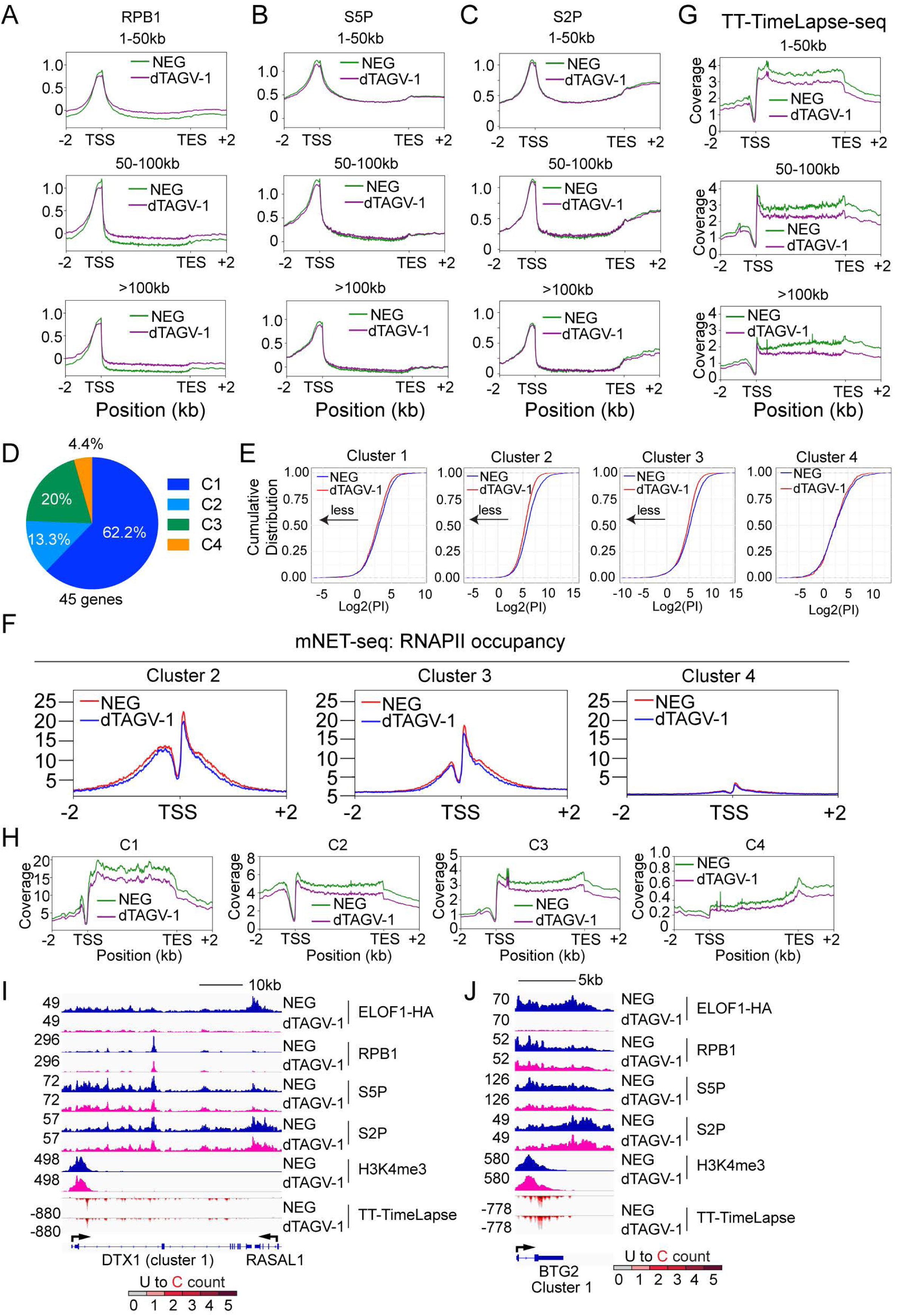
Related to Figure 4. (A-C) CUT&RUN-seq metagene plot of RPB1 (A), S5P-RNAPII (B), S2P-RNAPII (C) in genes separated into three groups by gene length. (D) The percentage of AID target genes in the four K-means clusters. (E) RNAPII pausing index (PI) in ELOF1-degron cells treated with NEG or dTAGV-1 for 4 hours followed by 2-day dox treatment. Cumulative index plots of pausing index were calculated from RPB1 CUT&RUN-seq data. (F) Metagene profiles aligned at the TSS showing mean occupancy of RNAPII (mNET-seq) upon ELOF1 degradation. Sense profiles are shown. (G) Metagene profile of TT-TimeLapse-seq in genes separated into three groups by gene length. (H) Metagene profile of TT-TimeLapse-seq in four clusters. (I, J) Genome browser tracks of ELOF1, RPB1, S5P-RNAPII, S2P-RNAPII, H3K4me3 CUT&RUN-seq and TT-TimeLapse-seq in *DTX1* (F) and *BTG2* (G) regions. Reads for TT-TimeLapse-seq categorized by U to C mutations.

**Figure S5.**
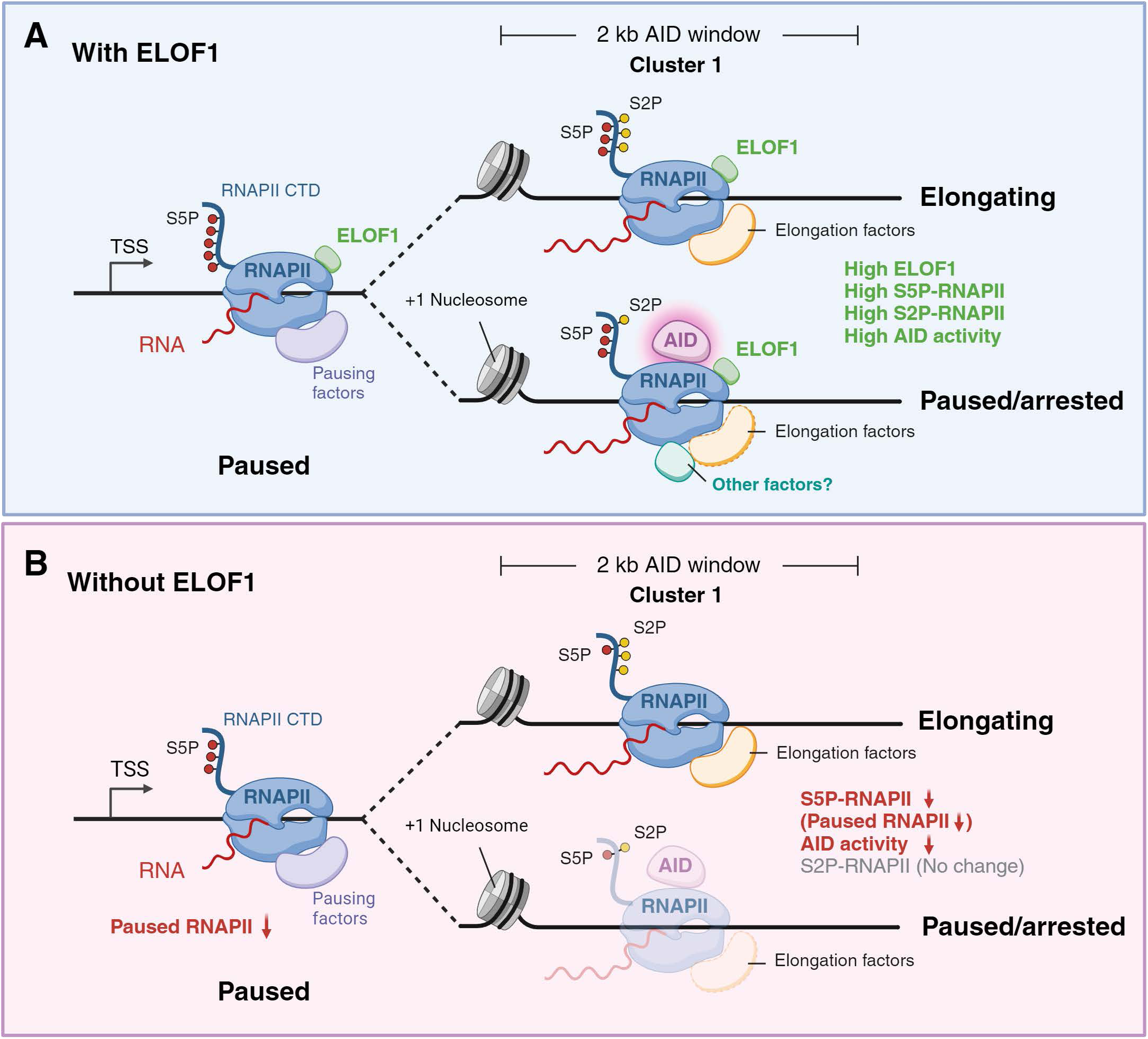
Model for role of ELOF1 in AID targeting. After transcription initiation, RNAPII typically pauses 25-50 bp downstream of the TSS and possesses high levels of serine 5 phosphorylation (S5P, red lollipops) of its RPB1 C-terminal domain (CTD; curved blue line). Release from the promoter-proximal pause site is accompanied by decreasing levels of S5P and increasing levels of CTD serine 2 phosphorylation (S2P, yellow lollipops). In the presence of ELOF1 (A), cluster 1 genes (which include most AID targets) are highly transcribed, acquire high levels of S2P, and maintain substantial levels of S5P in the 5’ portion of the gene. Most RNAPII complexes enter full elongation mode due to acquisition of elongation factors (upper line), while a small fraction become paused/arrested perhaps due to association with additional factors, creating the substrate for AID action (red halo) (lower line). In the absence of ELOF1 (B), levels of S5P-(but not S2P-) RNAPII are reduced throughout the gene, with a concomitant reduction in paused/arrested RNAPII substrate for AID and hence reduced AID-mediated DNA deamination. Reduced levels of S5P-RNAPII are depicted as being due to reduced levels of S5P phosphorylation per RNAPII complex but could also be due to reduced numbers of RNAPII complexes possessing the modification.

**Table S1. CRISPR loss of function screen data**

Genes are listed in rank order as positive acting SHM factors (sgRNAs enriched in GFP positive versus GFP negative cells), as computed by MAGeCK.

